# Characterization of the SARS-CoV-2 Spike in an Early Prefusion Conformation

**DOI:** 10.1101/2020.03.16.994152

**Authors:** Tingting Li, Qingbing Zheng, Hai Yu, Dinghui Wu, Wenhui Xue, Yuyun Zhang, Xiaofen Huang, Lizhi Zhou, Zhigang Zhang, Zhenghui Zha, Tingting Chen, Zhiping Wang, Jie Chen, Hui Sun, Tingting Deng, Yingbin Wang, Yixin Chen, Qinjian Zhao, Jun Zhang, Ying Gu, Shaowei Li, Ningshao Xia

## Abstract

Pandemic coronavirus disease 2019 (COVID-19) is caused by the emerging severe acute respiratory syndrome coronavirus 2 (SARS-CoV-2), for which there are no efficacious vaccines or therapeutics that are urgently needed. We expressed three versions of spike (S) proteins—receptor binding domain (RBD), S1 subunit and S ectodomain—in insect cells. RBD appears monomer in solutions, whereas S1 and S associate into homotrimer with substantial glycosylation. The three proteins confer excellent antigenicity with six convalescent COVID-19 patient sera. Cryo-electron microscopy (cryo-EM) analyses indicate that the SARS-CoV-2 S trimer dominate in a unique conformation distinguished from the classic prefusion conformation of coronaviruses by the upper S1 region at lower position ~15 Å proximal to viral membrane. Such conformation is proposed as an early prefusion state for the SARS-CoV-2 spike that may broaden the knowledge of coronavirus and facilitate vaccine development.

## Introduction

The novel coronavirus grouped in betacoronavirus genus has become the third serious virus intruder to human in the coronaviridae, after sever acute respiratory syndrome coronaviruses (SARS-CoV) and middle east respiratory syndrome coronavirus (MERS-CoV), recently named SARS-CoV-2. In the phylogenic tree of the coronaviruses, SARS-CoV-2 is genetically close to some bat coronavirus and SARS-CoV, however, with its origin undefined^1^. SARS-CoV-2 causative disease “Coronavirus disease 2019” (abbreviated “COVID-19”) is characterized by high fever, dry cough, difficulty breathing and sever atypical pneumonia, which usually be confirmed by virus RNA positive or pulmonary computed tomography (CT) in clinical practice^2, 3^. In terms of higher human-to-human transmissibility, SARS-CoV-2 has spread over 118 countries and areas, and led to over 125,288 confirmed cases worldwide and at least 4,614 deaths, as of March 12^th^ 2020. The World Health Organization (WHO) has declared the SARS-CoV-2 epidemic as a pandemic of international concern and updates the COVID-19 situation every day.

SARS-CoV-2 is an enveloped, single and positive-stranded RNA virus encapsulated with a genome of ~30 kb. At least three membrane proteins including the surface spike protein (S), an integral membrane protein (M), a membrane protein (E). Like other coronaviruses, S is responsible for initiating the engagement to a specific cellular receptor angiotensin-converting enzyme 2 (ACE2) and mediating the cell-virus membrane fusion by the class I fusion mechanism^4, 5^. Thus, S is the main target for neutralizing antibodies against viral infection and the core immunogen constituent of vaccine design. S is consisted of S1 and S2 subunits and the cleavage on S1/S2 boundary by protease during biosynthesis is prerequisite for coronaviruses cellular membrane fusion and subsequent infection^6^. SARS-CoV-2 evolves a 4-residue insertion (RRAR) as potential furin cleavage site rather than SARS-CoV and other bat coronaviruses, which may contribute to the higher transmissibility of this novel coronavirus^6, 7^. Previous studies suggested the infection process of MERS-CoV^8^ and SARS^9^ viruses, where S trimer undergoes conformational transition from a prefusion conformation ready for ACE2 binding to a postfusion conformation for eventual virus-cell membrane fusion. Structure determination of SARS-CoV and MERS-CoV spike trimers captured a variety of scenarios in the prefusion conformation showing partial (one or two) receptor-binding domain (RBD) in the “up” conformation and the rest in the “down”, and all in either “up” or “down”. The conformation transition from “down” to “up” could expose the receptor binding site, and the subsequent receptor engagement would lead to a substantial conformation rearrangement of S trimer from prefusion conformation to postfusion. Two recent studies^7, 10^ reported cryo-electron microscope (cryo-EM) structures of SARS-CoV-2 spike trimers in the prefusion conformation with 2 RBDs down and 1 RBD up. In the case of SARS-CoV, this conformational change during RBDs “down” to “up” was associated with the binding of receptor ACE2 as well as the recognition of neutralizing monoclonal antibodies ^11^.

A safe and efficacious vaccine is urgently needed to control and eliminate the SARS-CoV-2 infection. Various forms of vaccine candidates, mostly aiming to elicit neutralizing antibodies against S proteins, are under preclinical research or even subjected to clinical trials^12^. Here, we cloned S ectodomain and its fragments RBD and S1 into recombinant baculovirus and expressed the proteins in insect cells. We found that S and S1 formed homotrimer in solutions and the three proteins reacted well with COVID-19 convalescent human sera. Cryo-EM analysis demonstrated the S trimer unexpectedly retains a unique conformation distinguished from the classic prefusion conformation of coronavirus spikes, that may represent an early state rather than the known prefusion conformation of S spike. These results might broaden the knowledge on coronavirus virology and provide another protective conformation of S trimer for structure-based vaccine design against SARS-CoV-2 infection and its causative COVID-19.

## Results

### Construct design, expression and purification of SARS-CoV-2 S proteins

To screen a potent immunogen for COVID-19 vaccine development, we designed three constructs—S ectodomain, S1 and RBD—for the SARS-CoV-2 Spike (S) protein expression by aligning the SARS-CoV-2 S gene (Genbank accession no. NC_045512.2) to a SARS-CoV strain (Genbank accession no. NC_004718) S gene sequence in terms of structure-defined domain profile of the SARS-CoV S protein (Fig. 1A). The gene of SARS-CoV-2 S ectodomain encoding amino acids (aa) 15-1,213 with removal of its original signal sequence was cloned to the downstream of the gp67 signal sequence in pAcgp67B plasmid vector (Fig.1B). and with its C-terminal addition of a thrombin cleavage site, a T4 trimerization foldon motif and his tag. The segments S1 (aa 15-680) and RBD (aa 319-541) were cloned similar to S ectodomain, keeping gp67 signal peptide and his tag to facilitate secretory outside cell and affinity purification, respectively, but without the thrombin site and T4 foldon (Fig. 1A). The three constructed plasmids were respectively co-transfected into Sf9 insect cells with v-cath/chiA gene deficient baculovirus DNA for the generation and amplification of recombinant baculovirus, which were then harnessed to infect Hive Five insect cells to eventually produce recombinant proteins, named S, S1 and RBD respectively.

**Figure 1.**
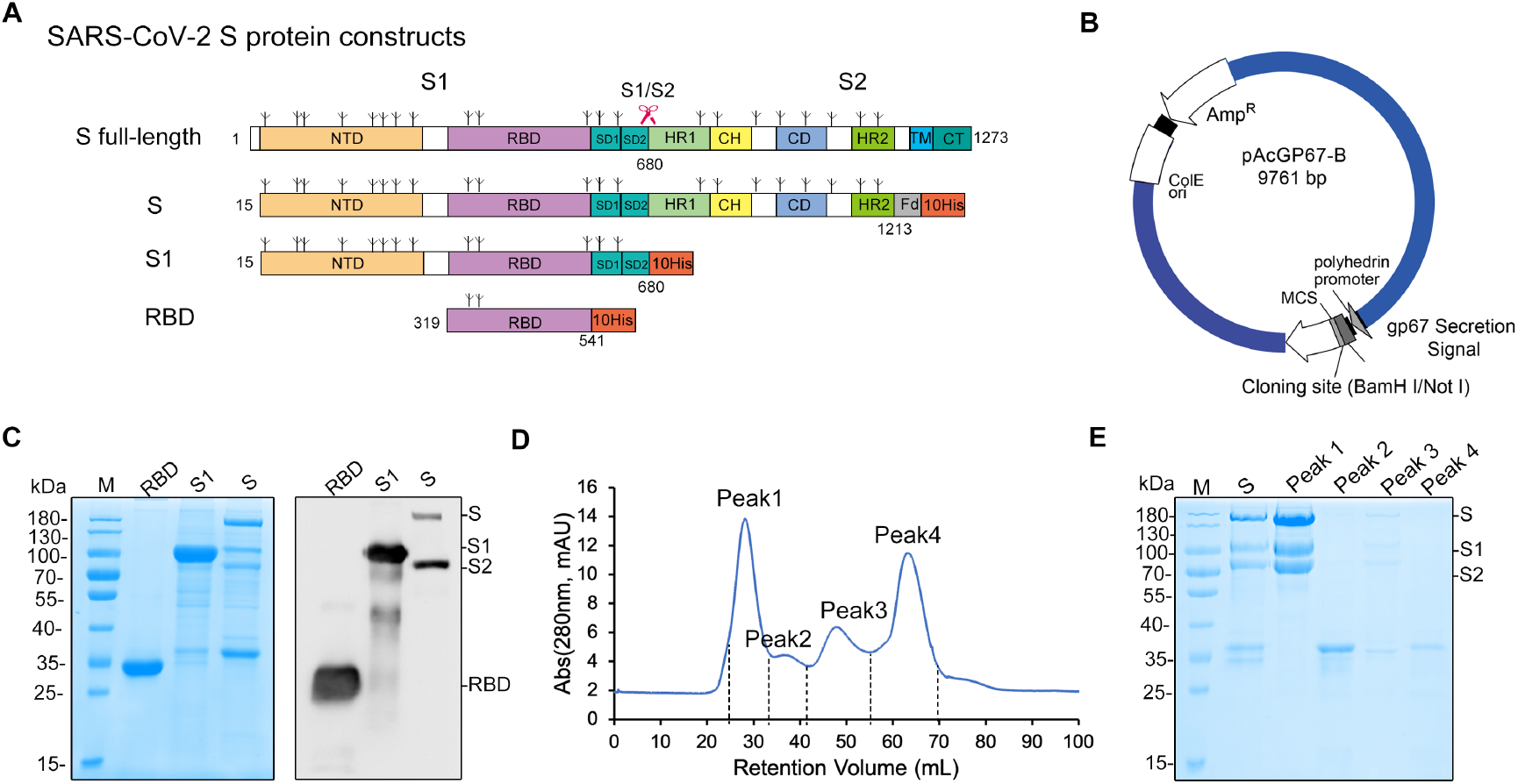
Schematic map of the SARS-CoV-2 S protein constructs. (A) Linear representations of the S protein primary structure and construct design. NTD, N-terminal domain; RBD, receptor binding domain; SD1, subdomain 1; SD2, subdomain 2; HR1, heptad repeat 1; CH, central helix; CD, connector domain; HR2, heptad repeat 2; TM, transmembrane domain; CT, cytoplasmic tail; FD, T4 foldon motif. The predicted glycosylation sites are indicated above the domain bars. (B) Map of the cloning vector pAcgp67B. The interest genes were cloned to plasmid pAcgp67B at BamH I/Not I site to generate transfer vectors. (C) SDS-PAGE and western blotting of the Ni-NTA purified proteins. RBS, S1 and S were eluted by 250 mM imidazole. Anti-His antibody was used as detection antibody in western blotting. (D) Size-exclusion chromatogram of the second-step purification of the S protein. (E) SDS-PAGE of the four fractions harvested from the chromatography purification as shown in (D).

The recombinant proteins were mostly soluble expressed and secreted into the culture medium. The centrifugation supernatants of cell culture went through metal affinity chromatography using Ni-NTA resin. S, S1 and RBD proteins were mainly eluted in a separation fractions under 250 mM imidazole elution, and resolved as molecular weight (m.w.) of ~180 kDa, 110 kDa and 35 kDa, respectively, in SDS-PAGE as indicated by a corresponding western blotting (WB) using anti-His antibody as detection antibody (Fig. 1C). Interestingly, about one half S proteins were cleaved into S1 (identical migration site to S1 lane in Fig. 1C) and S2 (about 80kDa developed in anti-His WB) possibly by innate furin of insect cell that was also found in other cases of enzymatic cleave while protein expression in insect cell, such as Flu HA ^13^. The eluted S fraction was further polished by Superdex 200 to remove contaminative proteins (Fig. 1D). These peaks fractionated at retention volume 28mL, 36mL, 48mL, and 65mL, were further harvested and subjected to SDS-PAGE analysis. The results indicated that S proteins together with cleaved S1/S2 were resolved at peak 1 in size-exclusion chromatography (Fig. 1D) and showed a high purity of over 95% total S/S1/S2 in gel (Fig. 1E). Overall, one-step Ni-NTA affinity chromatography produced RBD with 95% purity and a yield of 30 mg per L cell culture, S1 with about 90% purity and 10 mg per L yield, while further purification through a size-exclusion chromatography (SEC), the resultant S sample had over 95% purity regarding intact S and cleaved S1/S2, and was harvested in a yield of 1 mg per L cell culture. These data set up a start point for further optimization on expression and purification process of SARS-CoV-2 S immunogen candidates through insect baculovirus expression vector system (BEVS).

### Physiochemical properties of SARS-CoV-2 S-RBD, S1 and S proteins

We next investigated the physiochemical properties of the recombinant S protein and its fragments purified from insect cells, including association potential, thermal stability and glycosylation situation. Firstly, high pressure size-exclusion chromatography (HPSEC) and sedimentation velocity analytical ultracentrifugation (SV-AUC) analyses were carried out to measure the oligomerization potential of the three proteins in solution. RBD, S1 and S all showed single major peak in HPSEC profiles at elution volume of 9.0 mL, 5.5 and 5.3 mL, respectively (Fig. 2A and Fig. 2B). RBD, S1 and S were further verified by SV-AUC, where RBD sedimented as single species of 3.1S in c(s) profile, corresponding to apparent molecular weight 22 kDa (Fig. 2D); S1 existed as a dominant species of 11.3 S (estimated as 277 kDa corresponding to S1 trimer) and a minor aggregate form of 20 S (Fig. 2E); S and cleaved S1/S2 resolved as 15.2 S, equivalent to 577 kDa, approximately as the theoretical molecular weight of intact S trimer. The three proteins were further analyzed by differential scanning calorimetry (DSC) that was usually used to investigate the inner thermostability of macromolecules or their complexes^14^. RBD and S1 showed one major peak at comparable thermal denaturation midpoints (Tm) of 46.0 °C and 45.5 °C, respectively (Fig. 3G and 3H), whereas S sample showed two major peaks at Tm of 45.5°C (identical to Tm of S1) and 64.5°C (Fig. 3I), which might reflect the coexistence of intact S and cleaved S1/S2.

**Figure 2.**
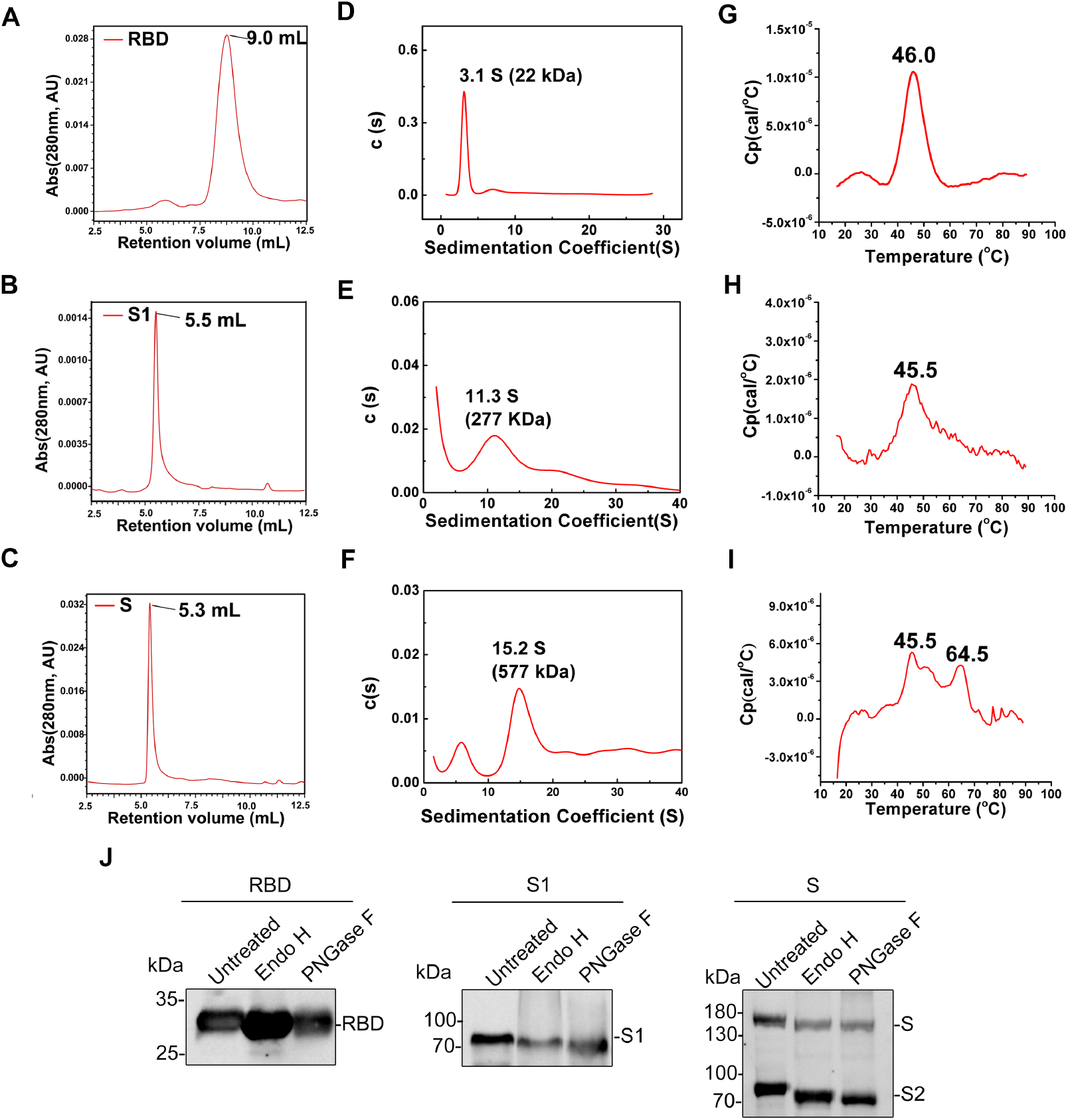
Characterization of the purified RBD, S1 and S proteins. (A-C) HPSEC profiles of the purified RBD, S1 and S proteins; (D-F) AUC profiles of RBD, S1 and S proteins; (G-I) DSC profiles of RBD, S1 and S proteins. (J) Western blotting of three purified proteins treated with Endo H and PNGase F or untreated as control. Anti-His antibody was used as detection antibody in western blotting.

**Figure 3.**
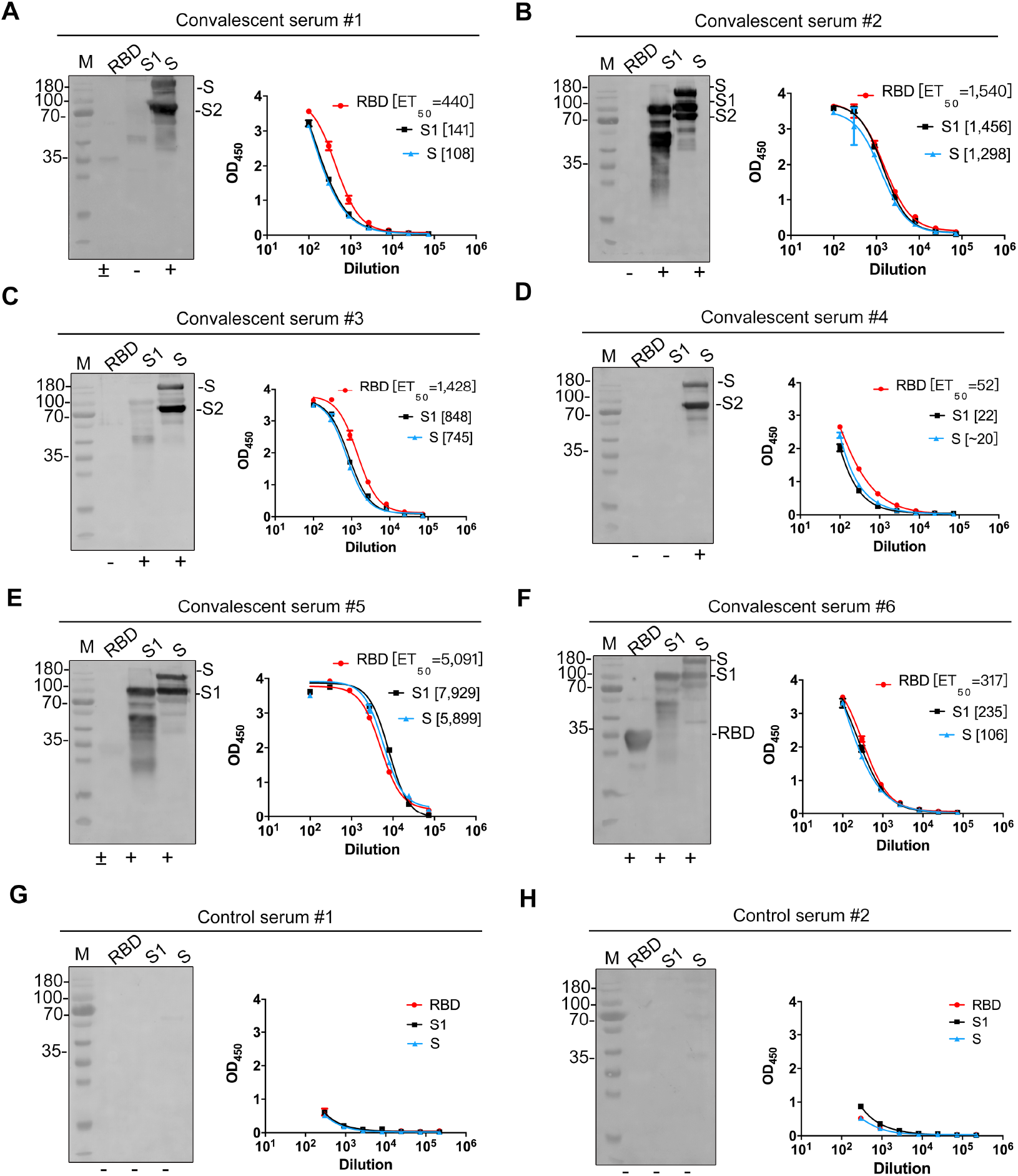
Antigenicity of RBD, S1 and S proteins against convalescent sera. (A-F) The reactivity of the RBD, S1 and S proteins against six COVID-19 convalescent human sera (#1-#6) by western blotting (left panel) and ELISA (Right panel). (G, H) Results of two control sera. The gels used for western blotting were duplicates of the reducing SDS gels same as Fig. 1C.

On the other hand, we investigated the glycosylation extent of the three protein by enzymatic deglycosylation analysis. Endo H could unleash the chithobiose core of high mannose and some hybrid oligosaccharides from N-liked glycoproteins, therefore remove the extended branches of glycans and leave the one N acetylglucosamine (GlcNAc) on N-linked glycoproteins. While PNGase F would release N-linked glycan moieties between GlcNAc and ASN residues within a glycoprotein. It should be noted that glycosylation in insect cells is featured as terminal mannose glycans, unlike complex sialylated glycans in mammalian cells, and glycosylation is known to correlate the immunogenicity and broad-coverage protection of a glycoprotein immunogen^15, 16^. After the treatment of either Endo H or PNGase F, RBD showed no discernible decrease of molecular weight in SDS-PAGE/anti-His WB, S1 and S2 both demonstrated nearly ~10 kDa decrease, and the intact S exhibited substantial shrinkage in molecular weight of about ~20 kDa decrease (Fig. 2J). The analyses conclude that the glycosylation extent within S glycoprotein is RBD < S1 ~ S2, consistent to the predicted glycosylation profile of S polypeptide (Fig. 1A).

### Reactivity of SARS-CoV-2 RBD, S1 and S proteins against convalescent COVID-19 human sera

We next evaluated the antigenicity of the three versions of S proteins by WB and ELISA using a panel of six COVID-19 convalescent human sera, which was collected from COVID-19 patients after they recovered from the disease in the First Affiliated Hospital of Xiamen University. Eight reducing SDS gel duplicates of the one depicted in Fig. 1C were prepared for WB analysis using these six convalescent sera and two control sera from health human (Fig. 3A-3H, left panel). As expected, intact S protein bands reacted well with all the six convalescent sera (Fig. 3A-3H, left panel). Unexpectedly, five of six sera showed no or very weak reactivities against RBD, only Serum #6 possessed RBD’s activity. Among the five sera with lower RBD-reactivity, Serum #2, #3 and #5 well recognized S1 and the cleaved S1 in lane S, suggesting these sera may specifically react with NTD of S1. S2 demonstrated reaction activity against all the six sera, like the intact S. No detectable reaction was observed in the control sera (Fig. 3G and 3H). Inconsistent to the WB results, RBD, S1 and S shared comparable reactivities against the convalescent sera in ELISA, although the sera per se presented varied reaction titers (represented as ET50) following the reaction sequence: Serum #5 > #2 > #3> #1 > # 6 > #4 (Fig. 3A-3H, right panel).

Taken together, RBD, S1 and S proteins from insect cells maintain the native-like SARS-CoV-2 epitopes. These epitopes in native virion should be immunogenic in COVID-19 patients and capable of eliciting high antibody titer in the convalescent phase of SARS-CoV-2 infection. Among these epitopes, most RBD epitopes are strictly tertiary conformation-dependent sites that are damaged upon the mild denatured condition of reductant and SDS treatment, NTD within S1 bears some linear epitopes, whereas S2 part essentially has linear epitopes that are immunogenic in all COVID-19 patients (n=6).

### Cryo-EM structures of SARS-CoV-2 S proteins

To examine the structure of the trimeric S ectodomain with native sequence, we prepared cryo-EM grids using the Ni-NTA purified S proteins and collected 1,513 electron micrograph movies. Most of motion-corrected micrographs demonstrated plenty of well-dispersed particles with an approximate size as the canonical coronavirus S trimer (Fig. 4A). A total of 162,645 particles were picked out for multiple rounds of 2D classification, consequently, 37,147 particles grouped into top 10 classes, rendering typical feature of S trimer in prefusion conformation as recently reported^7, 10^, were selected for further analysis (Fig. 4B). 3D reconstruction (applying 3-fold symmetry) yielded the density map of prefusion spike (S-pre) at resolution of 5.43 Å (Fig. 3D and Supplementary Fig. 1A).

**Figure 4.**
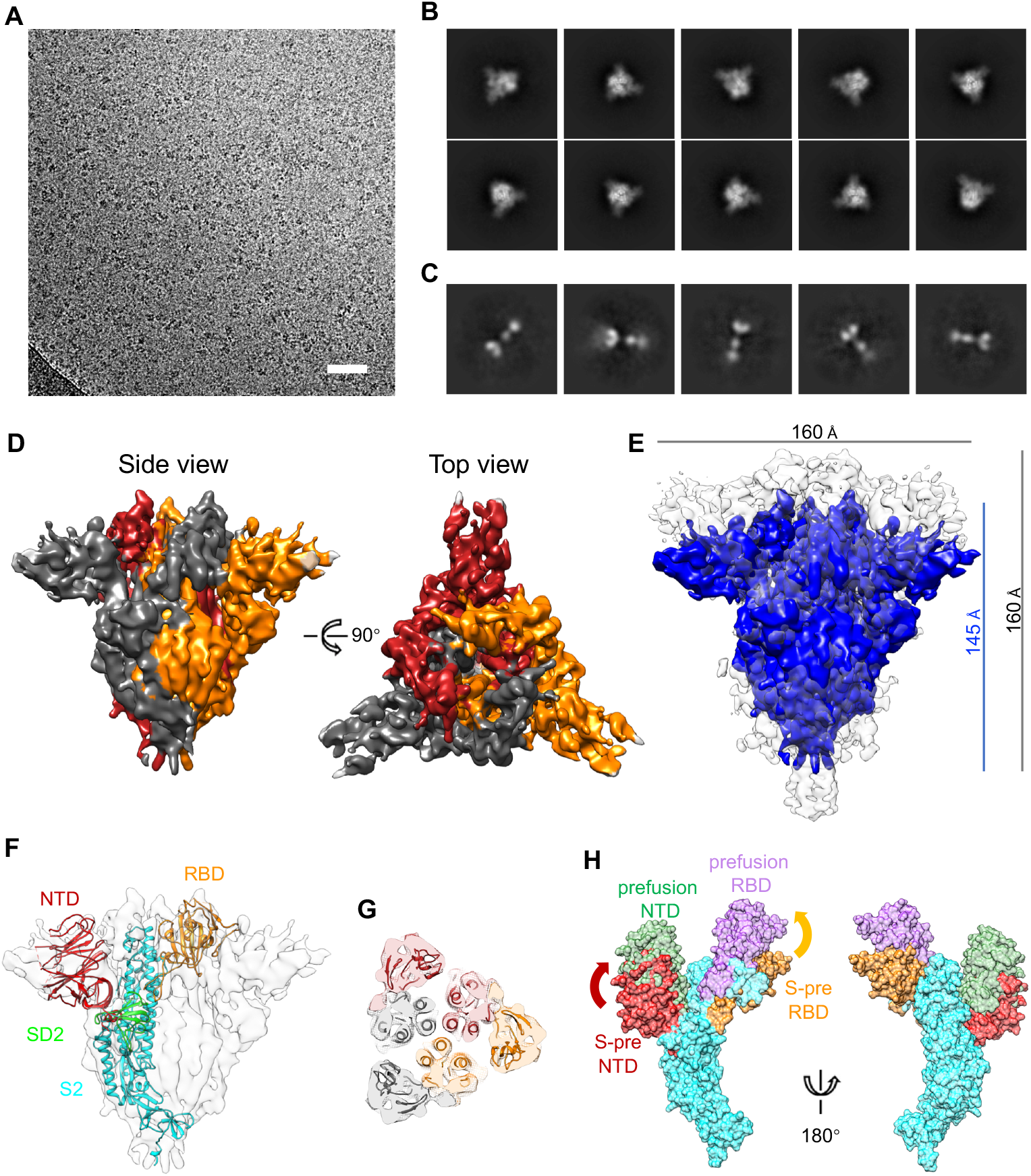
Cryo-EM structure of the SARS-CoV-2 S trimer. (A) Representative micrograph of frozen-hydrated SARS-CoV-2 S particles, Scale bar 50nm. (B, C) Ten and five selected class averages showing the particles along different orientations belonging to prefusion (B) and postfusion (C) S protein, respectively. (D) 5.43 Å density map of prefusion S trimer (S-pre) that is colored by protomer. (E) Structural comparison to the reported prefusion SARS-CoV-2 S trimer (EMD-21374, C3 symmetry, low-pass to 5.43Å) show a different conformation of S-pre (~15Å shorter in height). (F, G) Each domain of the model of prefusion SARS-CoV-2 S monomer (F) or trimer (G) (PDB no. 6VSB) were separately fitted in the density map of S-pre. (H) Schematic diagram shows conformational diversities between NTD and RBD of S-pre (NTD: red, RBD: orange) and reported prefusion S (NTD: light green, RBD: light purple).

Structurally, three S monomers intertwine around each other and associate to homotrimers with 145 Å height seen from side-view and 160 Å diameter in top-view (Fig. 4C and 4D). We then recruited the recently reported cryo-EM map of S prefusion trimer (EMD-21374, at resolution of 3.17 Å, low pass to 5.43 Å prior to structural comparison) and compared our cryo-EM map at same resolution (Fig. 4D). It was worthy noted that the compared prefusion SARS-CoV-2 S trimer was engineered with site-directed mutations to stabilize prefusion conformation and expressed in 239F cells. The mutant included two stabilizing proline mutations at residues 986, 987 and a “GSAS” substitution at the furin cleavage site^7^. Surprisingly, the alignment demonstrated that the two cryo-EM structures share similar mushroom-shaped architecture in particular nearly identical at stalk moiety (S2 region), but our S-pre shows the cap part (S1 region) at ~15Å lower position than the reported S trimer in RBD-down prefusion conformation (Fig. 4D). Regarding to substantial mismatch at the density of 3 S1 subunits, we respectively fitted 5 individual domains (NTD, RBD, SD1, SD2 and S2) of the SARS-CoV-2 S structure (PDB code 6VSB) to our S-pre map. In the fitting map, NTD, RBD, SD2 and S2 could be well placed in the S-pre map, especially for the latter two, which reflects the aforementioned good match at the stalk of the mushroom-shape (Supplementary Fig. 2). However, there is no observable density between RBD and SD2 to accommodate an SD1 model (Supplementary Fig. 2), which suggests SD1 region is dramatically flexible in our S-pre structure (Fig. 4E and 4F). When the combined model of fitted NTD-RBD-SD2-S2 was superimposed to the original S protomer structure (PDB code 6VSB, Chain A, RBD in down conformation), both NTD and RBD in the original S obviously move and rotate up against our combined model (Fig. 4G). The structural comparison demonstrated that the S-pre trimer retains a unique conformation different from the prefusion conformation of the two reported SARS-CoV-2 spike structures (PDB codes 6VSB and 6VXX).

We then compared the conformation of our S-pre structure with that of 21 deposited coronavirus S models. Six representative S structures^7, 17–20^ from four known genus (α-, β-, γ- and δ-genus) in coronaviridae are respectively fitted to the S-pre map (Supplementary Fig. 3). Five S trimer structures of other coronaviruses share similar prefusion conformation with the reported SARS-CoV-2 S structure but substantially distinct with the unique conformation of our S-pre.

Apart from most particles classified as S-pre in our sample, 2D classifications also showed five classes of few particles (2,951) assuming an elongated rosette-shape assembly. These particles were further reconstructed and yielded a structure at lower resolution of 8.40 Å (Supplementary Fig. 1B) that could be considered as post-fusion spike (S-post), as the structure has similar shape but shorter length (~170 Å) as compared to the postfusion spike of SARS-CoV^21^ and the presumed one observed in native SARS-CoV-2 virion (BioRxiv, https://doi.org/10.1101/2020.03.02.972927) (Supplementary Fig. 4). Fitting the S-post map with the core region structure of SAR-CoV-2 S2 subunit in post-fusion conformation (PDB code 6LXT) indicated that our S-post exhibits roughly rod shape similar with the post-fusion structure (Supplementary Fig. 4).

### Conformational transition of SARS-CoV-2 spike from early prefusion to postfusion

Briefly, we’ve obtained two conformations of SARS-CoV-2 spike from insect cells. The dominant one maintains the similar mushroom-shaped trimer as the previous models, while the S1 region substantially diverges. The other conformation essentially resembles the postfusion state. However, we could not find the classic prefusion conformation in our sample. We next tried to figure out at which stage the unique conformation occurs during the spike conformation change. The space relationship of NTD or RBD to S2 domain in the unique, RBD-down and RBD-up prefusion conformations (Fig. 5A) was measured by a reference plant approximately parallel to the viral membrane. The plane is defined by the positions of three equivalent Cα atoms (residue 694 be used) from three S2 subunits of the trimer structures. Numerical data shows that (1) the NTD and RBD in the unique conformation retain the lowest position in the three prefusion conformations; (2) the NTD and RBD of RBD-down prefusion stretch upward 16.6° rotation/16 Å elevation, and 13.1°/18 Å, respectively, with respect to the unique conformation; (3) from RBD-down to RBD-up prefusion state, the RBD elevates 9 Å with an additional rotation whereas the NTD remaining nearly stationary (Supplementary Movie S1). The resultant “up” RBD is ready for ACE2 binding and the spike eventually is rearranged to postfusion state upon RBD-ACE2 interaction^11, 22^ (Fig. 5B). The motion trend of NTD and RBD from prefusion to postfusion state in conformational transition is away from the viral membrane, suggesting the unique conformation may occur earlier than RBD-down prefusion conformation, named as “early prefusion conformation” (Fig. 5). This early prefusion conformation might exist in other coronaviruses as well.

**Figure 5.**
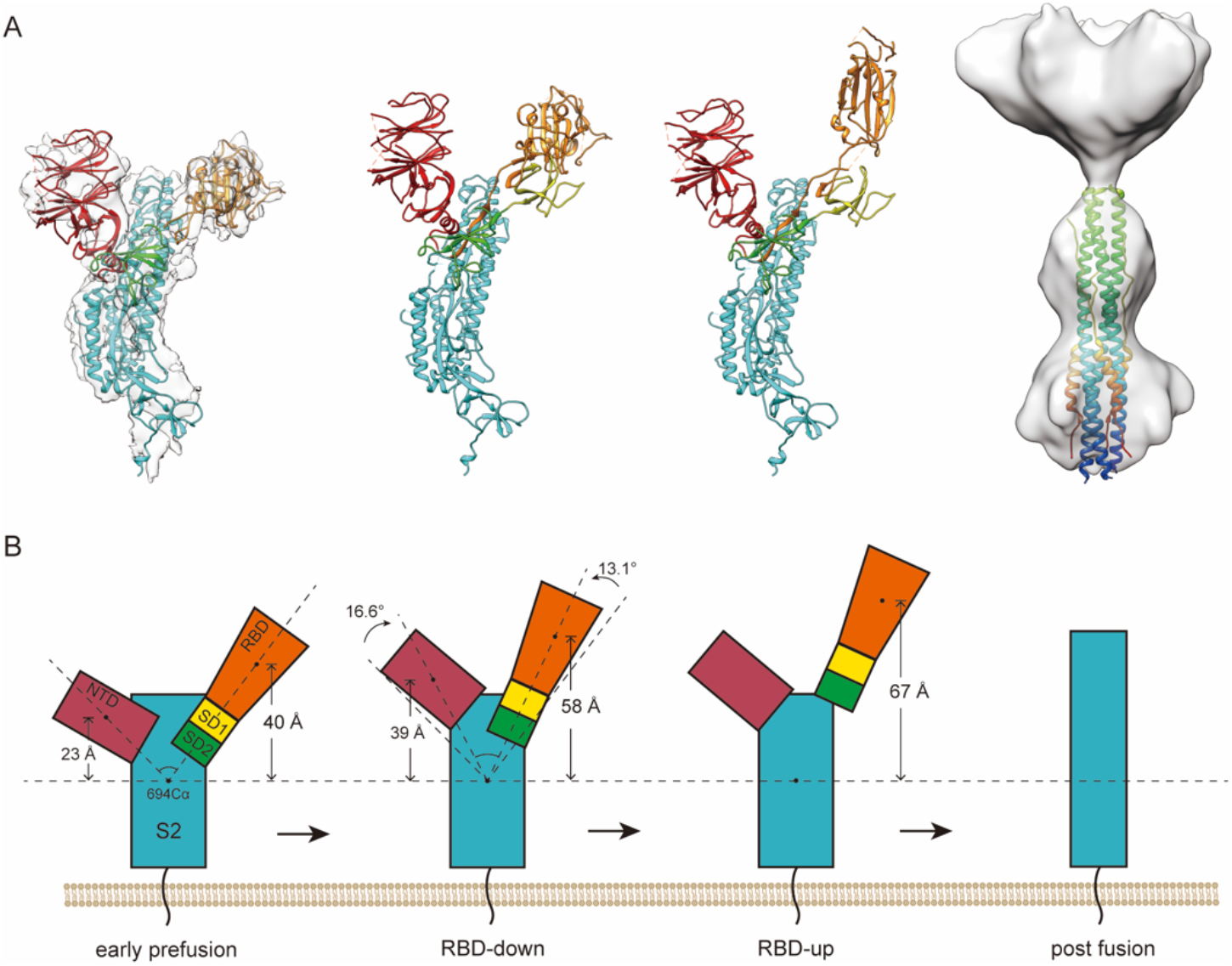
SARS-CoV-2 S spike at early pre-, pre- and post-fusion states and the proposed timeline for conformation change. (A) Four conformations of SARS-CoV-2 S spike. For clarity, only one monomer at early prefusion, RBD down prefusion and RBD up prefusion conformation is shown in ribbon mode. The model fitted in our S-pre map is a combined model comprised of individually fitted NTD, RBD, SD2 and S2 domain, which were evicted from the model of prefusion S trimer (PDB no. 6VSB). A deposited postfusion core of SARS-CoV-2 S2 subunit (PDB no. 6LXT) is fitted to our S-post map. Domains are designated by different colors, red: NTD, orange: RBD, yellow: SD1, green: SD2 and cyan: S2. (B) Simplified schematic diagram of the S monomer interpreting the conformation change across different states.

## Discussion

SARS-CoV-2 has crossed the species barrier and sweep over the planet by person-to-person transmission in an R0 ~2.56 rate^23^, first wave in China and the second wave booming outside China. WHO has declared the event as another pandemic infectious disease in human history, and the epidemiology of SARS-CoV-2 infection is still in data accumulation. Although the biology and virology of SARS-CoV-2 remain elusive, in terms of knowledge on other coronaviruses, the spikes decorating the SARS-CoV-2 virion play a critical role in viral attachment and entry to host cells. Cryo-EM structures of spikes in the prefusion conformation, and RBD-bound receptor ACE2 have indicated the engagement of SARS-CoV-2 to cellular membrane requires a serial of conformational change of RBDs. The change is presumed from the start point of 3 RBDs down in the prefusion conformation, then RBD(s) up for ACE2 binding, and eventually spike is rearranged to postfusion. In this study, we suggest that the SARS-CoV-2 spike may retain at more precedent state than the classic prefusion conformation that has been determined for other coronaviruses. This early prefusion conformation features that the cap of the mushroom-shaped spike constituted by three S1 subunits is more proximal to viral membrane by 15 Å than in the classic prefusion conformation.

The SARS-CoV-2 spike expressed in insect cells predominantly retains a unique early prefusion conformation, which was repeatable in at least three batches of samples and is ascribed to two possible reasons – native aa sequence used in the S ectodomain construct and over-expression in insect cells. There is about a half of S proteins undergoing cleavage on the S1/S2 boundary site in the purified samples both after the first Ni-NAT and the second SEC purification (Fig. 1C and 1E). Further analyses suggest the split between S1 and S2 likely takes no effect on the trimerization of S trimer. It is known that the insect cells can confer post-translation glycosylation for protein over-expression as mammalian cells despite the latter can produce more complex sialyation^24^, and thus provide an alternative way to generate glycoprotein in native conformation. Our results indicate that RBD, NTD and S2 domains of SARS-CoV-2 demonstrated different glycosylation extent in insect cells, however, RBD, S1 and S proteins comparably react well with six convalescent COVID-19 human sera albeit they differ in domain composition, polypeptide length and oligomerization.

There are numbers of SARS-CoV-2 vaccine candidates, including inactivated, vectored, recombinant and nucleotide vaccine forms, under preclinical research. Various versions of S proteins are the major targets for vaccine immunogen candidate. In addition to potent neutralizing antibody elicitation upon immunization, potential antibody-dependent disease enhancement (ADE) is the major concern for an efficacious SARS-CoV-2 vaccine. ADE has been found in the development of numbers of virus vaccine candidates, including respiratory syndrome virus (RSV), dengue fever ^25, 26^, human immunodeficient virus (HIV), SARS-CoV, MERS-CoV ^25–27^ and so on. It is believed that ADE is associated with non-neutralization epitope attribute and / or specific antibody isotype^28, 29^, in which virus-bound antibody would promote the viral infection to immune cells through Fc fragment targeting γFc receptors on the cellular surface and enhance the disease severity. Therefore, the strategy of vaccine design against SARS-CoV-2 should include the consideration of antigen region selection, glycosylation number/extent and exactly presented prefusion conformation. The prefusion conformation needed to be maintained is exemplified by the case of RSV vaccine candidate in which F trimer in prefusion is much potent than postfusion^30–32^. Hence the early prefusion conformation proposed for SARS-CoV-2 spike should be drawn an attention for immunogen design as well as the prefusion one.

In conclusion, we obtain three kinds of S proteins showing excellent antigenicity and find an early prefusion conformation for SARS-CoV-2 spike. Nevertheless, the molecular level detail for such conformation and the underlying immunogenicity should be further investigated, and whether this conformation recapitulates the exact state of spike in native SARS-CoV-2 virion remains to be determined.

## Materials and Methods

### Cloning, protein expression and purification

The SARS-CoV-2 S gene (Genbank accession no. NC_045512.2) was synthesized and cloned into a baculovirus shuttle vector pAcgp67B (BD Biosciences, CA, USA) using Gibson assembly. The S construct encoding aa 15-1,213 (numbered as original sequence), contains a thrombin site, a T4 foldon domain to assist in trimerization and a C-terminal 10-His tag for purification. For S1 construct contains gene encoding aa 15-680 followed by a 10-His tag. The RBD construct (aa 319-541) also contains 10-his tag to facilitate purification. In all three constructs, the natural signal peptide (aa 1-14 analyzed by SignalP tool) was replaced with a gp67 secretion signal peptide at N-terminus.

The expression and purification of proteins were performed as described previously^33^. All plasmids were co-transfected with linearized 2.0 DNA (deficient in *v-cath/chiA*genes) (Expression Systems, CA, USA) into *Sf9* insect cells (Thermo Fisher Scientific, MA, USA), according to the protocol provided by the manufacturer (Expression Systems). The transfection supernatant was harvested and amplified 2 times to obtain a high titer of the recombinant viruses. Hive Five cells (BTI-TN-5B1-4) (Thermo Fisher Scientific) were cultured in ESF921 medium (Expression Systems) and infected with recombinant virus at an multiplicity of infection (MOI) of 5 in the exponential growth phase (2× 106 cells/ml, 95% viability) at 28°C for 72 h. The culture media was centrifugated at 8,000 rpm for 20 min. Then the supernatant was dialyzed against phosphate-buffered saline (PBS), pH 7.4, and purified with Ni-sepharose fast flow 6 resin (GE Healthcare, Boston, USA) by the elution with 250 mM imidazole. The protein concentrations of the final purified samples were measured with Pierce™BCA Protein Assay Kit (Thermo Fisher Scientific).

### SDS-PAGE and western blot

Protein samples were mixed with loading buffer and boiled for 10 min, and subjected to sodium dodecyl sulfate-polyacrylamide gel electrophoresis (SDS-PAGE). Equal amounts of proteins for each sample were loaded onto two SDS-PAGE gels, one for western blotting and one for Coomassie staining. The proteins were electrophoresed for 70 min at 80 V in a BioRad MINI-PROTEAN Tetra system (BioRad Laboratories, CA, USA), and the gel was stained with Coomassie Brilliant Blue R-250 (Bio-Rad) for 30 min at room temperature. For western blotting, separated proteins were transferred onto a nitrocellulose membrane (Whatman, Dassel, Germany) using a Trans-Blot Turbo transfer system (Bio-Rad). The membrane was blocked and then incubated for 1 h with an His-tag-specific mouse mAb antibody (Proteintech, Rosemont, USA) or human sera (1:500 dilution). Unbound antibody was removed by five 5-min washes and the membrane was incubated with alkaline phosphatase-conjugated goat anti-mouse secondary antibody or goat anti-human IgG secondary antibody (Abcam, Cambridge, UK). Membranes were washed again and then developed using SuperSignal ELISA Pico Chemiluminescent Substrate Kit (Thermo Fisher Scientific).

### Enzyme-Linked Immunosorbent Assay (ELISA)

Purified proteins were coated onto 96-well microtiter plates at 100 ng/well in PBS at 37°C for 4 h. The background was blocked with 1 × Enzyme dilution buffer (PBS + 0.25% casein + 1% gelatin + 0.05% proclin-300) at 37°C for 2 h. Sera were diluted started at 1:100 followed with three-fold serially dilution, and added to the wells (100 μl) and incubated at 37°C for 1 h. Horseradish peroxidase (HRP)-labeled mouse anti-human antibody (Abcam) was used as secondary antibody at 1:5,000 for 30 min. Wells were washed again and the reaction catalyzed using o-phenylenediamine (OPD) substrate at 37°C for 10 min. The OD450nm (reference, OD620nm) was measured on a microplate reader (TECAN, Männedorf, Switzerland), with a cut-off value 0.1. The Half effective titers (ET50) was calculated by sigmoid trend fitting using GraphPad Prism software.

### Size-Exclusive Chromatography (SEC)

Ni-NTA purified S proteins were further loaded into Superdex200 (GE Healthcare), the fractions were harvested and analyzed by SDS-PAGE. All high-purity RBD, S1 and S proteins were subjected to HPLC (Waters; Milford, MA) analysis using a TSK Gel G5000PWXL7.8 × 300 mm column (TOSOH, Tokyo, Japan) equilibrated in PBS, pH 7.4. The system flow rate was maintained at 0.5 mL/min and eluted proteins were detected at 280 nm.

### Analytical Ultracentrifuge (AUC)

The AUC assay was performed using a Beckman XL-Analytical ultracentrifuge (Beckman Coulter, Fullerton, CA), as described elsewhere^34^. The sedimentation velocity (SV) was carried out at 20°C with diluted proteins in PBS. The AN-60 Ti rotor speed was set to 20,000-30,000 rpm according to the molecular weight of the control proteins. Data was collected using SEDFIT computer software, kindly provided by Dr. P. C. Shuck (NIH, Bethesda, MA, USA). Multiple curves were fit to calculate the sedimentation coefficient (S) using continuous sedimentation coefficient distribution model [c(s)], and then the c(s) used to estimate protein molar mass.

### Differential scanning calorimetry (DSC)

Differential scanning calorimetry (DSC) was carried out on the S proteins using a MicroCal VP-DSC instrument (GE Healthcare, MicroCal Products Group, Northampton, MA) as described previously^14^. In brief, all samples with a concentration of 0.2 mg/mL were measured at a heating rate of 1.5°C /min with the scan temperature ranging from 10°C to 90°C. The melting temperatures (Tm) were calculated using MicroCal Origin 7.0 (Origin-Lab Corp., Northampton, MA) software assuming a non-two-state unfolding model.

### Endo-H and PNGase-F digestion

The Endo-H (NEB) and PNGase-F (NEB) digestions were performed according to the protocol offered by instruction. In brief, the deglycosylation reactions were carried out using 10ug S proteins with 5uL of Endo H or PNGase F and incubated at 37°C overnights. The reactions were loaded in to SDS-PAGE and analyzed by Western blotting using anti-His as detecting anybody.

### Cryo-EM sample preparation and data collection

Aliquots (3 μL) of 0.5 mg/mL purified SARS-CoV-2 S protein were loaded onto glow-discharged (60 s at 20 mA) holey carbon Quantifoil grids (R1.2/1.3, 200 mesh, Quantifoil Micro Tools) using a Vitrobot Mark IV (ThermoFisher Scientific) at 100% humidity and 4°C. Data were acquired using the EPU software to control a FEI Tecnai F30 transmission electron microscope (ThermoFisher Scientific) operated at 300 kV. and equipped with a ThermoFisher Falcon-3 direct detector. Images were recorded in the 58-frame movie mode at a nominal magnification of 93,000X with a pixel size of 1.12 Å. The total electron dose was set to 46 e^−^ Å^−2^ and the exposure time was 1.5 s. 537 micrographs were collected with a defocus range comprised between 1.5 and 2.8 μm.

### Cryo-EM data processing

Movie frame alignment and contrast transfer function estimation of each aligned micrograph were carried out with the programs Motioncor^35^ and Gctf^36^. Particles were picked by the ‘Templete picker’ session of cryoSPARC v2^37^. Two rounds of reference-free 2D classification were performed and well-defined particle images were selected and non-uniform 3D refinement, 3D reconstruction with C3 symmetry were performed using cryoSPARC v2. The resolutions of the final maps were estimated on the basis of the gold-standard FSC curve with a cutoff at 0.143^38^. Density-map-based visualization and segmentation were performed with Chimera^39^.

## Acknowledgments

This work was supported by grants from the National Natural Science Foundation (grant no. U1705283, 31670935, 81971932, 81991491, 31730029), and the Major Project of Fujian Provincial Science Foundation for COVID-19 Research (Grants 2020YZ014001).

## Author Contributions

Y.G, S.L. and N.X. designed the study. T.L., Q.Zheng., H.Y., D.W., W.X., Y.Z., X.H., L.Z., Z.Zhang., Z.Zhai., T.C., Z.W., J.C., H.S. and T.D. performed experiments. T.L., Q.Z., H.Y., Y.W., Y.C., Q.Zhao., J.Z., Y.G., S.L. and N.X. analyzed data. T.L., Q.Z., H.Y., Y.G., and S.L. wrote the manuscript. T.L., Q.Zheng., H.Y., D.W., W.X., Q.Zhao., J.Z. Y.G., S.L., and N.X. participated in discussion and interpretation of the results. All authors contributed to experimental design.

## Supplementary Information

**Supplementary Figure 1**

**Supplementary Figure 2**

**Supplementary Figure 3**

**Supplementary Figure 4**

**Supplementary Movie 1**

