## Supplementary information for "Characterization of the SARS-CoV-2 Spike in an Early Prefusion Conformation"

Tingting Li et al.

**Supplementary Figure 1**

**Supplementary Figure 2**

**Supplementary Figure 3**

**Supplementary Figure 4**

**Supplementary Movie 1**

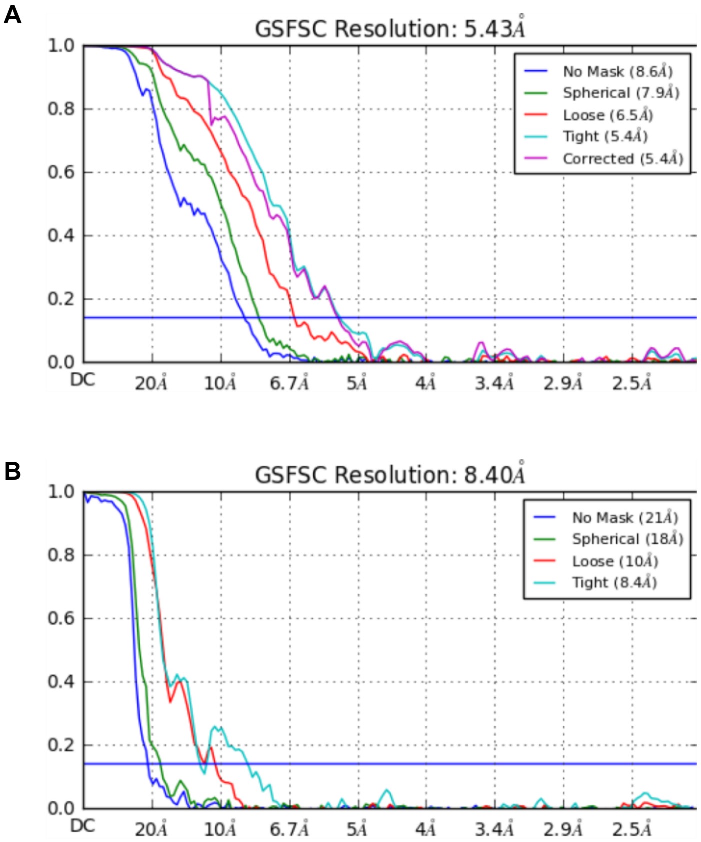


**Supplementary Figure 1. FSC curves of 3D reconstructions of S-pre (A) and S-post (B).**

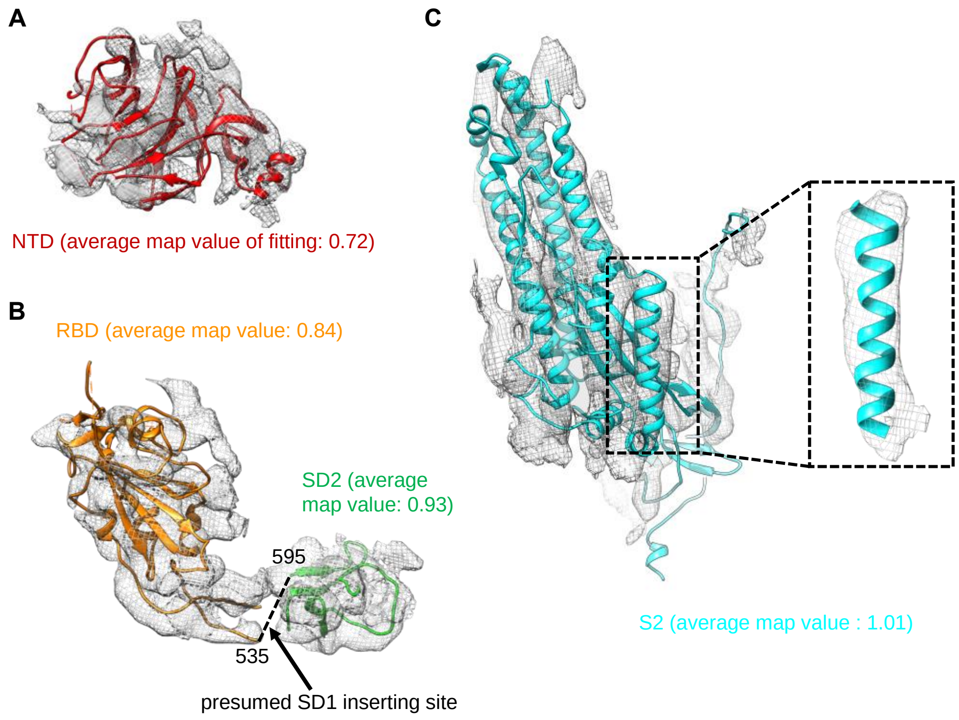


**Supplementary Figure 2. Fitting of NTD (A), RBD (B, left), SD2 (B, right) domains and S2 subunit (C) into segmented density maps from S-pre structure.** Average map values (generated by "fitting" tool in Chimera) of each domain were shown. The density of SD1 located between RBD and SD2 is smeared in our S-pre map as shown in (B).

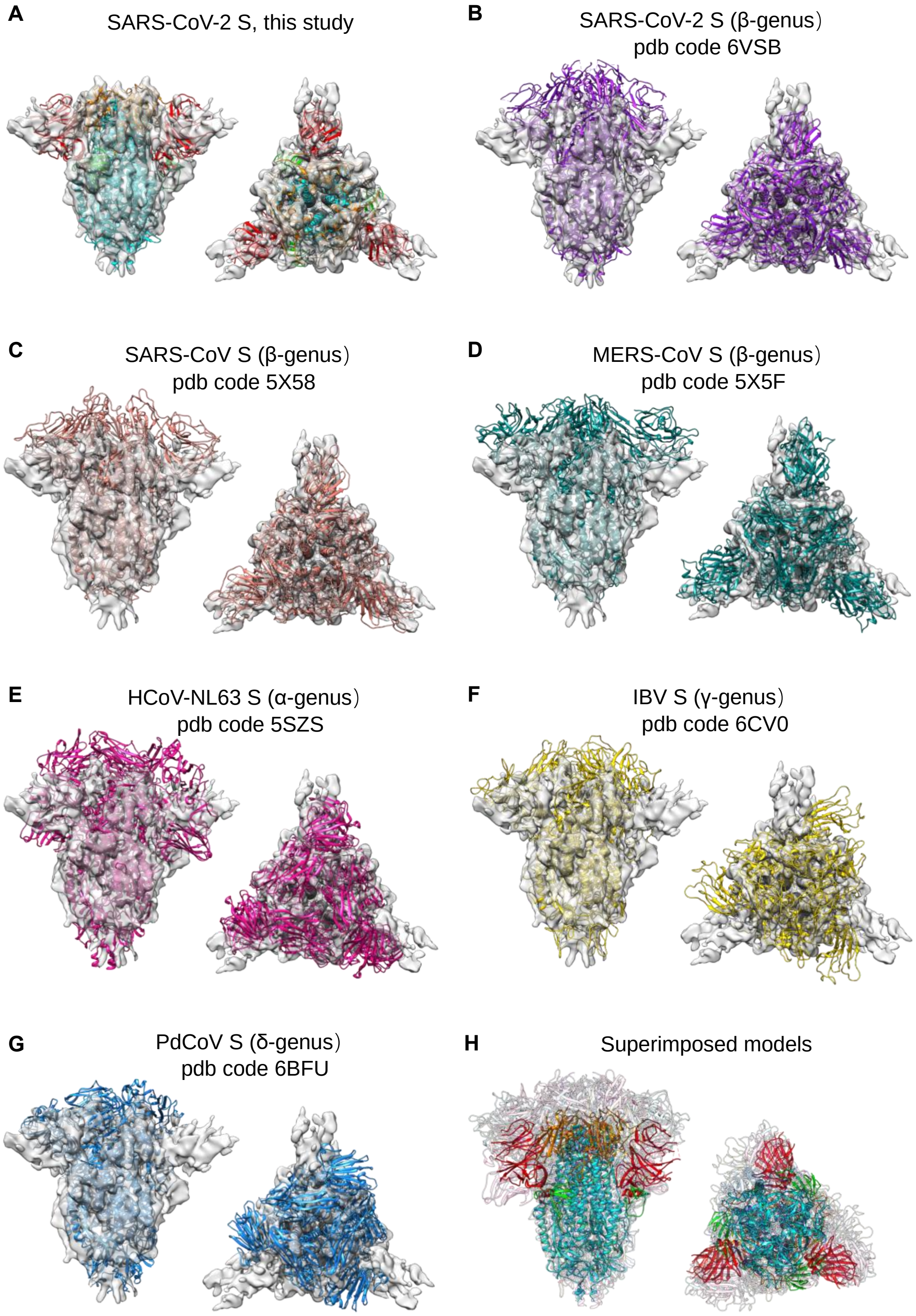


**Supplementary Figure 3. Structural comparisons S-pre of SARS-CoV-2 to prefusion S trimers from six representative coronaviruses.** A. density map and fitted model of S-pre. (B-G) Six representative S trimer structures from 4 genus of coronavirues were fitted respectively in the density map of S-pre. (H) superposition of above seven structures.

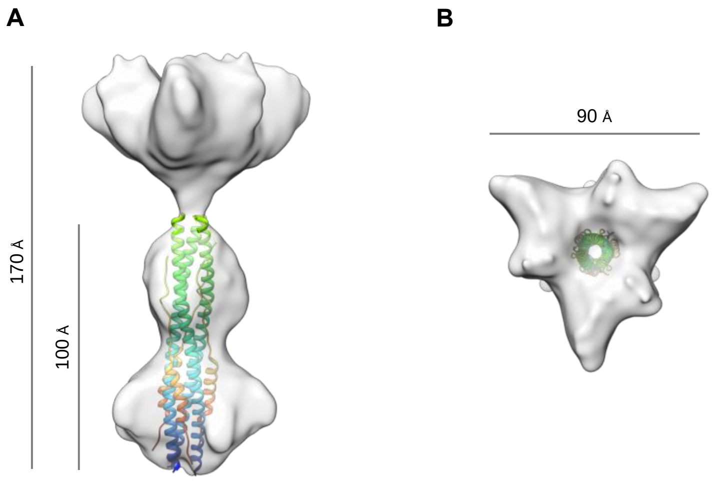


**Supplementary Figure 4. Cryo-EM structure of S-post at resolution of 8.40 Å.** Side (A) and top (B) view of S-post are shown, with 170Å in height and 90Å in diameter, respectively. A postfusion core of SARS-CoV-2 S2 subunit (PDB no. 6LXT) was fitted in the rod region of S-post (about 100Å in height).
